# The Porcine Skin Microbiome Exhibits Broad Fungal Antagonism

**DOI:** 10.1101/2024.01.12.574414

**Authors:** Karinda F. De La Cruz, Elizabeth C. Townsend, JZ Alex Cheong, Rauf Salamzade, Aiping Liu, Shelby Sandstrom, Evelin Davila, Lynda Huang, Kayla H. Xu, Sherrie Y. Wu, Jennifer J. Meudt, Dhanansayan Shanmuganayagam, Angela LF Gibson, Lindsay R Kalan

## Abstract

The skin and its microbiome function to protect the host from pathogen colonization and environmental stressors. In this study, using the Wisconsin Miniature Swine™ model, we characterize the porcine skin fungal and bacterial microbiomes, identify bacterial isolates displaying antifungal activity, and use whole-genome sequencing to identify biosynthetic gene clusters encoding for secondary metabolites that may be responsible for the antagonistic effects on fungi. Through this comprehensive approach of paired microbiome sequencing with culturomics, we report the discovery of novel species of *Corynebacterium* and *Rothia*. Further, this study represents the first comprehensive evaluation of the porcine skin mycobiome and the evaluation of bacterial-fungal interactions on this surface. Several diverse bacterial isolates exhibit potent antifungal properties against fungal pathogens *in vitro*. Genomic analysis of inhibitory species revealed a diverse repertoire of uncharacterized biosynthetic gene clusters suggesting a reservoir of novel chemical and biological diversity. Collectively, the porcine skin microbiome represents a potential unique source of novel antifungals.

**Highlights:**

- Porcine skin bacterial communities are consistent with previous reports on porcine and human skin.
- Fungal community composition resembles mycobiomes from other mammalian skin, but not human skin.
- Bacteria isolated from porcine skin have antimicrobial and particularly strong antifungal activity *in vitro*.
- Discovered three new *Corynebacterium* species and one new *Rothia* species.

## 1. Introduction

In 2019, an estimated 4.95 million people died globally due to complications of antimicrobial resistant infections (Murray et al., 2022). An additional 2 million annual deaths are attributed to fungal pathogens (McDermott, 2022). Many of these pathogenic fungi have developed resistance to at least one class of antifungals, making the need for novel antifungals even more dire. In recent years there has been increased attention to exploring naturally occurring competition between microbes within animal microbiomes to spur the discovery of novel antimicrobials (Cruz et al., 2022; Miethke et al., 2021). Due to the high similarity to human skin and overlap of the skin microbiome, we sought to characterize the bacterial and fungal communities of porcine skin. Using a combination of high-throughput sequencing and culturomics based approaches, we then evaluated bacterial isolates for antimicrobial activity against human pathogens, including a phylogenetically diverse panel of fungal species. Insights into antagonistic interactions within the porcine skin microbiome support the development of next-generation antimicrobial therapeutics.

Porcine and human skin are highly similar in structure (Summerfield et al., 2015). Both have thick epidermal and dermal layers that are firmly attached to the underlying subcutaneous connective tissue and sparse hair (Summerfield et al., 2015). These features make pigs a great corollary model for studying human skin function and wound healing. Pig and human skin bacterial community compositions are highly similar at the phyla and genera levels (Wareham- Mathiassen et al., 2023). At the phyla level skin communities are dominated by *Bacillota*, *Bacteroidota*, *Actinomycetota*, and *Pseudomonadota* and 97% of the genera are shared.

However, key differences exist in the specific bacterial species that reside on human versus porcine skin (Wareham-Mathiassen et al., 2023). These subtle species-level differences may provide untapped resources for identifying microbial-microbial interactions and characterizing the molecules that drive them. Moreover, the fungal communities within the pig skin microbiome and their interactions with the surrounding bacteria have yet to be characterized.

Biosynthetic gene clusters (BGCs) are grouped genes within a genome that encode the biosynthesis of secondary metabolites, which often benefit the microorganism and sometimes the host (Medema et al., 2015; Xia et al., 2022). Production of these secondary metabolites can serve to maintain their niche within microbiomes through securing nutrients and inhibiting the growth of potential competitors or pathogens. For example, microorganisms within porcine gut microbial communities utilize BGCs that inhibit the growth of gastrointestinal pathogens, which ultimately helps maintain community homeostasis and benefits the host (Wang et al., 2023). However, the BGCs within the porcine skin microbiome and the metabolites they produce have yet to be explored. By characterizing these BGCs, we aim to discover novel antimicrobials for use against prominent and emerging pathogens.

This work aims to characterize the bacterial and fungal communities on porcine skin and then identify isolates for potential antimicrobial activity against human pathogens. This was accomplished by high-throughput sequencing of the bacterial 16S rRNA gene and fungal internal transcribed spacer 1 (ITS1) in microbiome samples collected from Wisconsin Miniature Swine™ (WMS™). Each sample was also processed to maximize the diversity of isolates cultured. Isolates were then evaluated for *in vitro* activity antimicrobial against a panel of human bacterial and fungal pathogens, and their BGCs were explored for predicted antimicrobial activity based on whole genome sequences. Here we present i) characterization of the bacterial and fungal communities of the porcine skin and feces; ii) identification of bacteria able to inhibit the growth of human pathogens through secreted metabolites; iii) identification of novel species of *Corynebacterium* and *Rothia*; and iv) genomic characterization of BGCs predicted to have antifungal activity. To our knowledge, this is the first study to characterize the mycobiome and investigate BGCs encoding for antimicrobials within the porcine skin microbiome.

## 2. Methods

### 2.1 Sample collection

Samples were collected from two 6-8 month old male WMS™ pigs bred and maintained at the University of Wisconsin - Madison. Swabs for microbiome analysis were collected at five time points over two weeks. Swabs from the dorsal and ventral skin regions were collected after saturating in sterile 1X phosphate-buffered saline (PBS). The dorsal swabs were taken from the paraspinal, caudal region, while the ventral swabs were collected from the abdomen close to the midline. Fecal swabs were collected by inserting a sterile swab 1.0-1.5 inches through the rectal sphincter and gently rotating. The swabs for DNA extraction were stored at -20°C, prior to being sent to the UW Biotechnology Center for processing. Samples for the culturing and isolation of microbiota were also collected from each animal from six body sites, including the dorsal and paraspinal skin, the ventral abdomen (close to midline), and the inguinal, the axilla, oral, and nares regions. All swabs for microbial culture were stored at 4°C for up to 4 hours before being processed.

### 2.2 DNA extraction, library construction, sequencing

DNA extraction on samples collected from the dorsal skin, ventral skin, and feces was performed by the UW Biotechnology Center following the protocol for the DNAeasy 96 PowerSoil Pro QIAcube HT Kit (Qiagen, Hilden, Germany). Bacterial 16S rRNA gene amplicon libraries targeting the V3-V4 region or the fungal nuclear ribosomal internal transcribed spacer (ITS) regions ITS1-ITS2 were constructed using a dual-indexing method and sequenced on a MiSeq platform with a 2x300 bp run format (Illumina, San Diego, CA) at the University of Wisconsin – Madison Biotechnology Center. Air swabs taken roughly two to three feet from the pig served as the negative controls. ZymoBiomics Microbial Community Standard (Zymo Research, Irvine, CA) served as a positive control.

### 2.3 Culturing and bacterial isolation

Culture swabs were diluted to 1x10^-1^ in 1X PBS and 100µl was plated on Tryptic Soy Agar (TSA) with 5% sheep blood (BBL, Sparks, MD), brain heart infusion (BHI) agar (Millipore Sigma, Darmstadt, Germany), and BHI agar with 10% Tween 80 (VWR, Radnor, PA). Plates were incubated at 35°C overnight. To isolate culturable bacteria, colonies with distinct morphology were isolated on the respective plates the colony grew on, incubated at 35°C overnight. Isolated colonies were grown in BHI with 10% Tween 80 broth overnight at 35°C in a shaking incubator. To identify each bacterial isolate, a small portion was used for polymerase chain reaction (PCR) amplification and Sanger sequencing. The remaining isolate liquid culture was stored in 10% glycerol at -80°C.

### 2.4 Colony polymerase chain reaction and isolate identification

To create a PCR master mix for each isolate that was undergoing PCR, 12.5 µL of EconoTaq PLUS green 2X Master Mix (VWR, Radnor, PA), 1 µL of 10 µM 16S 8F primer, 1 µL of 10 µM 16S 1492R primer, and 10 µL PCR grade nuclease-free water (VWR, Radnor, PA) were combined. An aliquot (24.5 µL) of the master mix was added to all wells of a 96-well PCR plate (VWR, Radnor, PA). Then, 0.5 µL of overnight liquid culture was added to one well and repeated for each liquid culture. PCR was performed using a Veriti 96-Well Fast Thermal Cycler with the following parameters set: initial denaturation at 95°C for 10 minutes, 35 cycles of denaturing at 95°C for 30 seconds, annealing for 54°C for 30 seconds, and extension at 72°C for one minute, ending with a final extension at 72°C for five minutes. The plates were sent to the UW Biotechnology Center for Sanger sequencing of the bacterial 16S rRNA gene using the standard 27F primer on a 3730xl Genetic Analyzer.

### 2.5 Bioassays

All bacterial isolates were grown on BHI with 10% Tween 80 agar from glycerol stocks stored at -80°C. After growing at 37°C overnight, a single colony was picked and inoculated in 3 mL BHI with 10% Tween 80 broth to grow in a shaking incubator at 37°C overnight. Bacterial and fungal pathogens were grown under the same conditions. 24-well plates were filled with 1.5 mL basic media (BM) agar. Five µL of one bacterial isolate liquid culture was swabbed on the left side of each well in the plate. This was repeated for each isolate. The plates were incubated at 37°C until half the well was filled with the target isolate. The overnight pathogen liquid cultures were diluted 1:10 with 1X PBS. Three µL of one diluted pathogen liquid culture was pipetted to the right half or culture-free side of each experimental plate in one well. This was repeated for each pathogen in a well uninoculated by another pathogen. A 24-well plate with BM agar was inoculated with diluted pathogen liquid culture to serve as a control plate to compare to the experimental plates. The plates were incubated again for up to four days at 37°C to allow the pathogens to grow which was assessed by checking if the control plate fully grew. The ability of the skin bacterial isolates to inhibit the growth of the pathogens was assessed by a 0-3 scoring system: 0 – no inhibition; 1 – slowed or reduced growth; 2 –inhibition of the pathogen in a semi- circle pattern; 3 – total inhibition of the pathogen. Refer to **(Fig. S1)** for an example of the scoring system and infographic on the bioassay protocol.

### 2.6 DNA extraction and whole genome sequencing

Twenty-five bacterial isolates were selected for whole genome sequencing. BHI with 10% Tween80 agar plates was used to streak out the desired bacteria for DNA extraction from the freezer stock stored in 10% glycerol at -80°C. Cells were harvested directly from the plate and suspended in 300µL 1X PBS. DNA extraction was performed as described by the GenElute Bacterial Genomic DNA Kit (Sigma-Aldrich, Darmstadt, Germany) using the kit’s columns and reagents following the protocol for Gram-positive bacteria for all samples. The concentration of DNA was measured utilizing the Qubit 4 Fluorometer (ThermoFisher Scientific, Waltham, MA) and accompanying protocol for dsDNA with high sensitivity. Collection tubes were stored at - 20°C until sent for sequencing. Extracted DNA concentrations measuring above 3.3 ng/µL were sent to SeqCoast Genomics (Portsmouth, NH, USA) to perform short-read whole genome sequencing.

### 2.7 Sequence analysis

The QIIME2 (Bolyen et al., 2019) environment was used to process bacterial 16S rRNA gene sequencing and fungal ITS sequencing data. For the 16S rRNA gene data, paired-end reads were trimmed, quality filtered, and merged into amplicon sequence variants (ASVs) using DADA2 (Callahan et al., 2016). Taxonomy was assigned using a naive-Bayes classifier pre-trained on full-length 16S rRNA gene 99% OTU reference sequenced based on the SILVA SSU database (release 138). For the ITS gene data, single-end forward reads were trimmed and quality-filtered into amplicon sequence variants (ASVs) using DADA2. Taxonomy was assigned using a naive-Bayes classifier pre-trained on full-length eukaryotic nuclear ribosomal ITS gene 99% OTU reference sequenced based on the UNITE database (Version 8.0 2020-04-02).

### 2.8 Microbiome and statistical analysis

Data was imported into RStudio (Version 2023.03.1+446) running R (version 4.2.3), using the qiime2R package (v0.99) and analyzed using the phyloseq package (Bisanz, 2018; Davis et al., 2018; McMurdie and Holmes, 2013). ASVs were manually removed from sample data if they i) were identified as contaminants within the negative controls ii) were not identified at the phylum or genus level, iii) were determined to be eukaryotic, archaeal, mitochondria, or chloroplasts sequences, or iv) were an ASV below a 5% prevalence threshold. The ggplot2 and gplots packages were used to build plots (Galili et al., 2020; Wickham, 2016). The vegan package (version 2.6-5.0) was used to analyze microbial alpha diversity with the Shannon index and beta diversity via the weighted UniFrac metric (Oksanen et al., 2020). Differences in microbial beta diversity between groups were evaluated with type-2 permutation ANOVA (type-2 PERMANOVA). The MaAsLin2 package was used to analyze the differential relative abundance of specific taxa between various sample groups (Mallick et al., 2021). Further statistical analyses were conducted in R Studio running R.

### 2.9 Whole-genome assembly and taxonomy assignment

Whole-genome Illumina sequencing reads were processed and assembled as previously described (Salamzade et al., 2023a). Briefly, sequencing reads were processed for adapters using TrimGalore (v0.6.5)(Krueger et al., 2023), and trailing poly G-tail artifacts were trimmed using fastp (v0.21.0)(S. Chen et al., 2018). Next, genomic assembly was performed using Unicycler (v0.4.6) with default settings (Wick et al., 2017). Bacterial genomes were taxonomically classified using GTDB-tk (v2.1.1) with GTDB release 207 (Chaumeil et al., 2020; Parks et al., 2022) **(Supplementary Table S1)**. Genomes were further assessed for completeness and contamination using CheckM2 (Chklovski et al., 2023) **(Supplementary Table S1).** To assess the number of distinct novel *Corynebacterium* species, FastANI (Jain et al., 2018) was used to estimate the average nucleotide identity (ANI) between pairs of genomes lacking a species-level designation by GTDB-tk. Single-linkage clustering was used to group isolates into three unique species clusters based on pairs exhibiting ≥95% ANI (Olm et al., 2020). FastANI was further used to validate that these novel *Corynebacterium* species had < 95% ANI to genomes in the Skin Microbial Genome Collection, which was recently released and reported the discovery of multiple novel *Corynebacterium* species (Saheb Kashaf et al., 2021). To understand the phylogenetic placement of novel *Corynebacterium* species within the genus, a phylogenomic model was constructed of the *Corynebacterium* genomes in this study and representative genomes for the genus based on 95% ANI dereplication, as described previously (Olm et al., 2017; Salamzade et al., 2022). Phylogenomics was performed using GToTree with single copy core gene sets for Actinomycetota (Lee, 2019). The resulting phylogeny was visualized using iTol (Letunic and Bork, 2019).

### 2.10 Identification and analysis of biosynthetic gene clusters

The web-interface for antiSMASH (v7.0.0) (Blin et al., 2023) was used for the identification of BGCs in each isolate’s genome using the “relaxed” mode with all extra features or analyses requested. BiG-SCAPE (v1.1.5) (Navarro-Muñoz et al., 2020) was nextrun with the options “-- mibig --mix --include_singletons” to group BGCs across the genomes into gene cluster families (GCFs). To investigate the novelty of BGCs from newly discovered species, the BiG-FAM webserver (Kautsar et al., 2021) was used.

### 2.11 Data availability

Sequence reads for this project can be found under NCBI BioProject PRJNA1055114. Code for analysis and generation of figures can be found on GitHub at https://github.com/Kalan-Lab/DeLaCruz_Townsend_etal_PorcineSkinMicrobiome

## 3. Results

### 3.1 The porcine skin and fecal fungal mycobiome

To assess the porcine mycobiome, swabs of skin and fecal microbial communities underwent high-throughput sequencing of the fungal internal transcribed spacer (ITS1) of the fungal ribosomal RNA cistron. Across skin and fecal communities, *Basidomycota* and *Ascomycota* were the most abundant phyla **(Fig. 1A).** *Cutaneotrichosporon* was the most abundant genera on dorsal skin from one porcine subject while *Apiotrichum* was the most abundant genera in a second animal **(Fig 1B).** *Aureobasidium* and *Fusarium* were the most abundant on the ventral skin communities of pigs 1 and 2, respectively. *Trichosporon* and *Torulaspora* dominated fecal communities in porcine subjects. The differential relative abundance of fungal taxa was evaluated via MAASLIN2 (Mallick et al., 2021). Notably, Ascomycota was significantly more abundant in the fecal communities of pig 2 (FDR corrected p-value < 0.2, **Fig. S2A**), demonstrating inter-individual variability of the porcine mycobiome.

**Figure 1.**
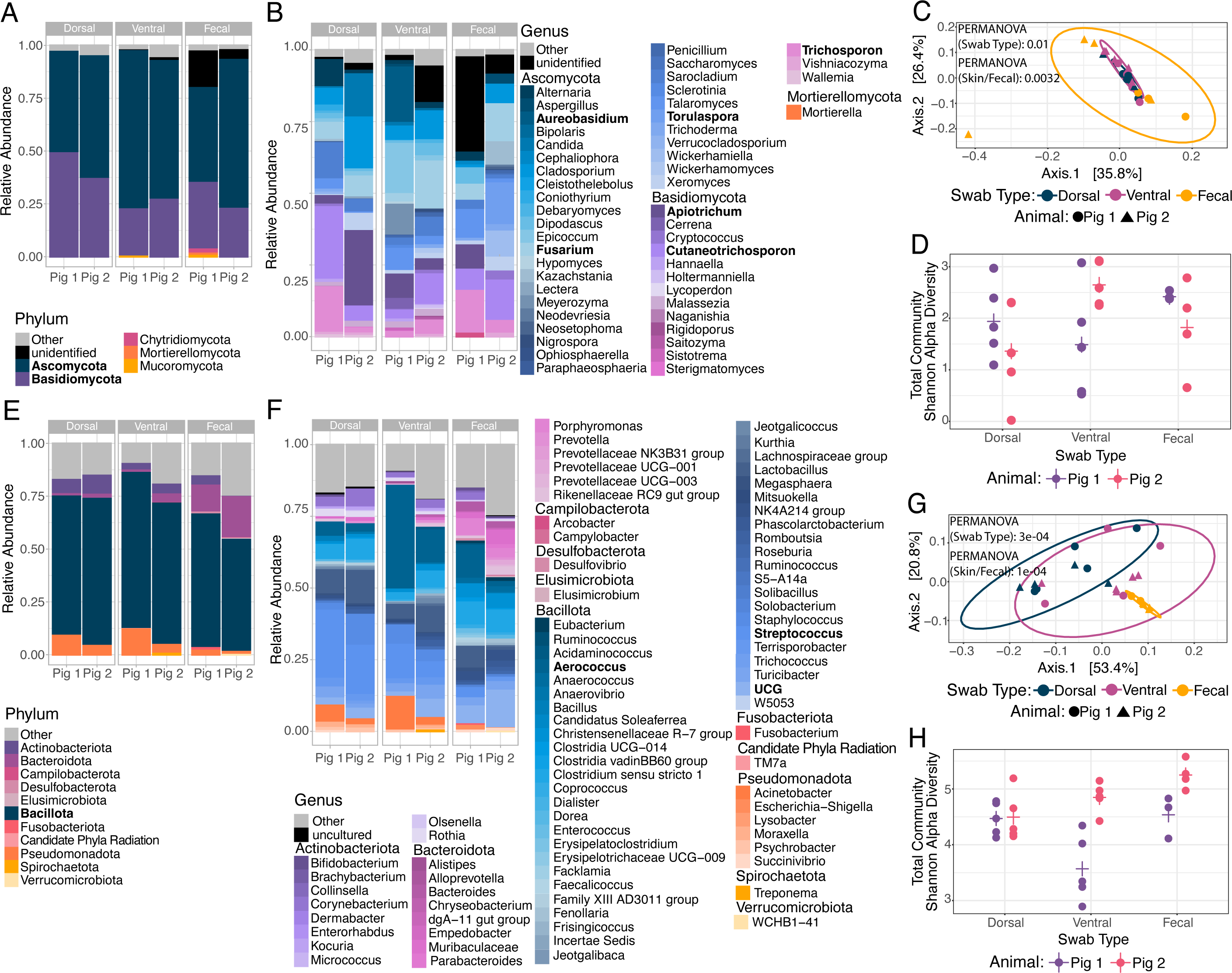
**Fungal and bacterial community compositions on porcine skin and feces**. Swabs of from dorsal and ventral skin sites as well as feces underwent high-throughput sequencing of the bacterial 16S ribosomal RNA gene and fungal ITS1 region. **A-B:** Relative abundances plots for skin and fecal mycobiomes at the phyla and genera level respectively. Bolded fungal names indicate taxa present in the greatest relative abundance at the skin or fecal sites for one of the animals. **C:** A weighted UniFrac beta diversity PCoA plot demonstrating the similarity of skin and fecal fungal community compositions. Groups compared via type-2 PERMANOVAs. **D:** Fungal community Shannon alpha diversity for each pig at each of the sites. **E-F:** Relative abundance plots for skin and decal microbiomes at the phyla and genera level respectively. Bolded bacterial names indicate taxa present in the greatest relative abundance at the skin or fecal sites for one of the animals. **G:** PCoA plot for the weighted UniFrac beta diversity of skin and fecal bacterial communities. Groups compared via type-2 PERMANOVAs. **H:** Bacterial community Shannon alpha diversity within the dorsal, ventral, and fecal communities.

Weighted UniFrac beta diversity was utilized to evaluate the compositional similarity of the porcine skin and fecal communities. This revealed that the dorsal skin, ventral skin, and fecal fungal communities are significantly distinct from one another (p-value < 0.05, type-2 Permutation ANOVA, **Fig. 1C**). Combined dorsal and ventral skin versus fecal fungal community compositions were also significantly different from one another (p-value < 0.01, type-2 PERMANOVA, **Fig. 1C**). Diversity of fungal taxa within each sample (alpha diversity) was evaluated with the Shannon index. There were no significant differences in fungal taxa alpha diversity between the subjects or across the skin and fecal sites (all p > 0.05, Mann-Whitney t-test. **Fig. 1D**). Collectively, these findings suggest that fungal communities within porcine feces and on porcine skin are more similar than their respective bacterial communities (discussed below).

### 3.2 Porcine skin sites and feces have distinct bacterial communities

To evaluate bacterial microbial communities on porcine skin and in feces, samples underwent high-throughput sequencing of the V3-V4 region of the bacterial 16S rRNA gene. Overall, the most abundant phylum across both skin and fecal samples was *Bacillota* **(Fig. 1E).** Compared to skin microbial communities, phyla unique to the fecal microbiome include *Desulfobacteriota*, *Elusimicrobiota*, *Fusobacteriota*, and *Verrucomicrobiota*. *Candidate Phyla Radiation* was unique to the skin sampling sites. Skin microbial communities were dominated by taxa within the *Streptococcus* and *Aerococcus* genera **(Fig. 1F).** The dominant genera within fecal communities differed by pig, with *Aerococcus* and *UCG* dominating the communities of one animal. Within these communities, dorsal skin had greater abundance of taxa within the Actinobacteriota phyla, *Clostridia* UCG-014, and *Moraxella* genera compared to ventral skin (FDR corrected p-value < 0.3, **Fig. S2B-D).** More detail on the differential abundances of skin and fecal taxa between each pig can be found in **Supplemental Figure S2E-I**.

The similarity of the porcine skin and fecal microbial communities was evaluated using the weighted UniFrac beta diversity metric. Fecal microbial communities were similar between porcine subjects, with *Aerococcus* and UCG in the highest relative abundance (**Fig. 1F-G**). Overall, bacterial community compositions of skin sites are distinct from fecal communities (p-value < 0.001, type-2 PERMANOVA, **Fig. 1G**) and bacterial communities between skin sites are also distinct from one another (p-value < 0.001, type-2 PERMANOVA, **Fig. 1G**).

Significant differences in the Shannon alpha diversity of samples between animals and across samples from the dorsal skin, ventral skin, or feces were observed (**Fig. 1H**). Bacterial diversity differed between animals, where pig 1 had lower alpha diversity than pig 2 for both the ventral skin site and fecal samples. Bacterial communities were also more diverse than fungal communities across skin sites and feces (16S v. ITS Shannon alpha diversity 4.15 ± 0.35 v. 1.65 ± 0.86 on dorsal skin, p-value < 0.01; 3.97 ± 0.75 v. 2.07 ± 0.97 on ventral skin, p-value < 0.01; 4.67 ± 0.42 v 2.08± 0.71, p-value < 0.05 in fecal communities. Wilcoxon matched-pairs single rank tests).

### 3.3 Bacteria isolated from porcine skin inhibit bacterial and fungal pathogens in vitro

To evaluate potential fungal-bacterial interactions occurring within the porcine skin microbiome we obtained representative bacterial isolates from the porcine skin microbiome using culture-based methods. Swabs were collected from the nares, dorsal, inguinal, oral, axilla, and ventral body sites for each porcine subject. Each swab was processed by culturing on multiple types of media to select for the highest phylogenetic diversity (see methods). Bacterial isolates were identified through Sanger sequencing of the full-length 16S rRNA gene. The majority of taxa isolated across all six sampled sites were within the *Actinobacteriota* (**Fig. 2A**). Species within the *Corynebacterium* genera were the most frequently isolated from the nare, inguinal, axilla, and ventral body sites, while *Rothia* species were most frequently isolated from dorsal skin, and *Kocuria* species were the most frequently isolated from oral samples (**Fig. 2B**).

**Figure 2.**
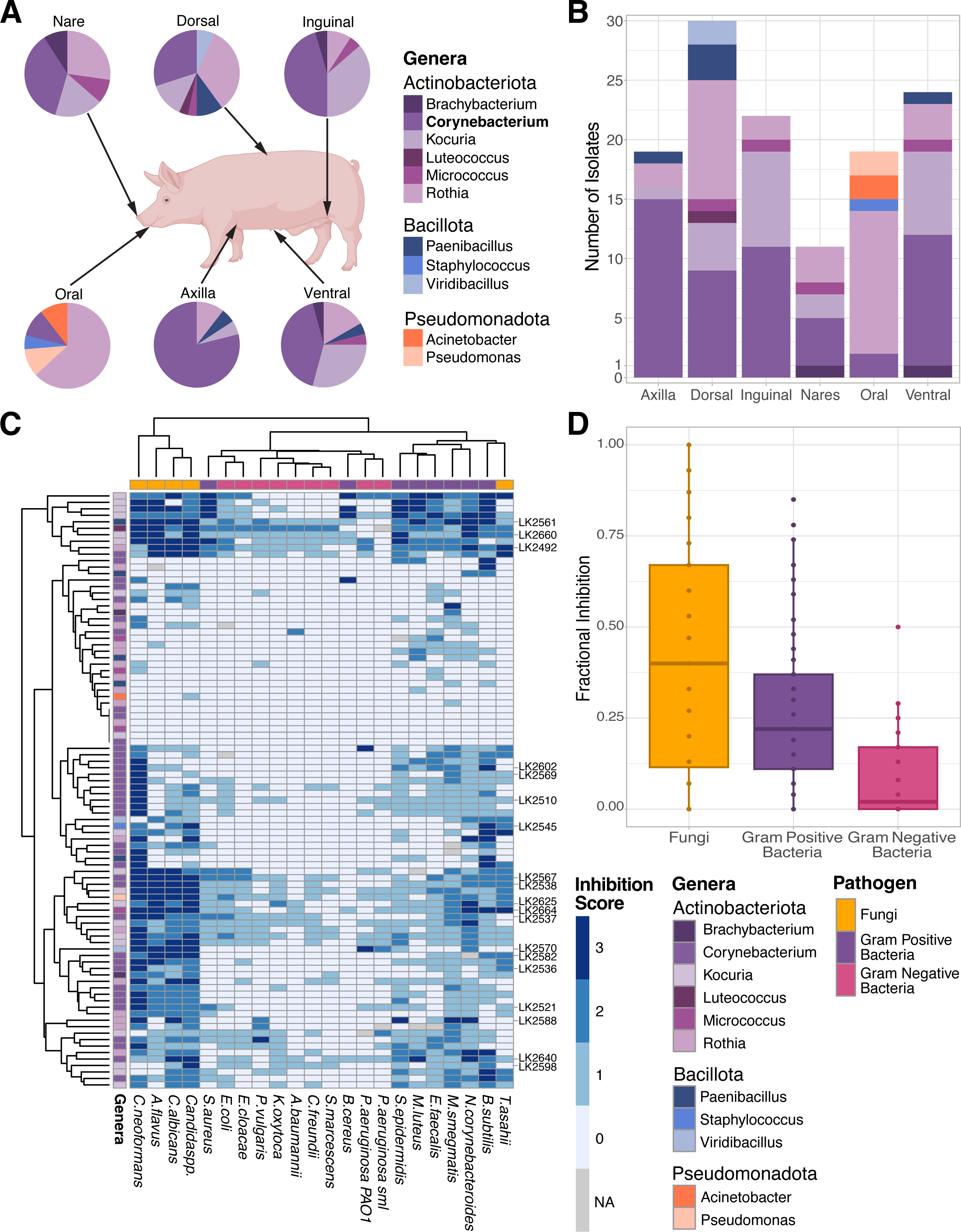
Bacteria isolated from porcine skin inhibit the growth of human pathogens. Bacteria were cultured and isolated from six porcine skin sites, including the skin surrounding the nares, oral, axilla, dorsal, ventral, and inguinal regions. All isolated bacteria were identified via Sanger Sequencing of the full length bacterial 16S rRNA gene. **A:** displays the relative frequency each taxa was isolated from each body site. The bolded genus indicates the genera isolated the most frequently. **B:** The total number of isolates, colored by genera, collected from each body site. The colors for each genera in plot A are the same for plot B. **C:** Isolated bacteria were evaluated for anti-microbial activity against a panel of human pathogens. **Supplemental Figure S1** details the methods for this assessment. This heat map details the inhibition scores for each isolate against the panel of pathogens. A score of 0 indicates that the isolate did not display antimicrobial activity, the pathogen had uninterrupted growth; 1 indicates slowed growth of the pathogen; 2 indicates the isolate significantly inhibited, but did not fully eliminate pathogen growth (eg. ring or small growth around inoculation spot); and a score of 3 denotes complete inhibition of pathogen growth. Isolates that underwent further evaluation with whole genome sequencing are indicated on the right-hand side of the heat map. **D**: A fractional inhibition was calculated on each of the isolates to generate a normalized score between zero and one for each group of pathogens (fungi, gram positive bacteria, gram negative bacteria). Zero indicates the bacterial isolate did not inhibit any of the pathogens in the group. One indicates complete inhibition of all pathogens in that pathogen group. Details on the fractional inhibition scores from each of the isolates tested are in **Supplemental Table S2.**

A direct coculture bioassay was then employed to evaluate the ability of skin-associated bacteria to inhibit the growth of known pathogens, including phylogenetically diverse fungal pathogens (**Fig. S1**). We found that out of the 22 pathogens evaluated, fungal pathogens, including *Cryptococcus neoformans*, *Aspergillus flavus*, *Candida albicans*, *Candida spp.*, and *Trichosporon asahii* were more frequently inhibited by isolates from porcine skin (**Fig. 2C**). The six bacterial pathogens that were most often inhibited by these isolates were *Nocardia corynebacteroides*, *Mycobacterium smegmatis*, *Enterococcus faecalis*, *Micrococcus luteus*, and *Staphylococcus epidermidis* (**Fig. 2C**). Overall, skin isolates within the genera, *Viridibacillus*, *Rothia*, *Corynebacterium*, and *Kocuria*, were responsible for the strongest inhibition of pathogens (**Fig. 2C**). Fractional inhibition values were calculated to provide standardized values to compare the overall antifungal and antibacterial activities of each isolate. A score of zero indicates the pathogens in the group were not inhibited in the presence of the isolated bacteria and a score of one indicates complete inhibition of all pathogens in that pathogen group. All bacteria isolated from the porcine microbiome, except for taxa within the *Paenibacillus* genera, displayed stronger inhibition of fungal pathogens compared to Gram-positive and Gram-negative bacterial pathogens (**Fig. 2D**). Details of the fractional inhibition values of each isolate are provided in **Supplemental Table 2**. Collectively, our findings exemplify that bacteria from the porcine skin can prevent the growth of bacterial and particularly fungal pathogens, which may mirror the bacterial-bacterial and bacterial-fungal interactions that routinely occur to maintain homeostasis of the skin microbial community, hallmarked by low fungal diversity.

### 3.4 Cultivation of bacterial isolates and genomics-based characterization of their biosynthetic potential uncovers novel species and BGCs

To explore the potential mechanisms bacteria utilize to inhibit fungi, we sequenced 25 bacterial isolates with strong antimicrobial activity from six different porcine body sites and characterized their biosynthetic gene clusters (BGCs). Each isolate genome was taxonomically classified using GTDB. From the 25 isolates that were sequenced and taxonomically classified, 14 (56%) were determined to belong to one of four novel species, including three distinct species of *Corynebacterium* (**Fig. 3**). Investigations of the placement of these novel species within the genus of *Corynebacterium* using phylogenomics, revealed that two of the species, indicated in blue (species 1) and green (species 2) in **figure 3**, were very closely related genetically (**Tables S1 & S2**). Using bioassay profiles in combination with genetic similarity estimates, we determined that our 25 isolates represented 19 unique strains **(Fig. 4A, Tables S1 & S2).** Of these, 10 were *Corynebacterium,* three were *Kocuria*, two were *Rothia*, and the remaining four were singletons belonging to the genera of *Micrococcus*, *Viridibacillus*, *Paenibacillus*, and *Staphylococcus* (**Fig. 4A**). Most of the isolates were obtained from the dorsal and ventral body sites. Analysis of the BGCs found within the genomes of these isolates revealed higher BGC diversity on the dorsal region with 34 different gene cluster families (GCFs). This finding is likely attributed to the greatest number of distinct genera being cultivated from the site (**Fig. 4A-B, Fig S3, Table S3**). Collectively, these findings indicate that the porcine skin microbial communities contain diverse BGCs across different body sites.

**Figure 3.**
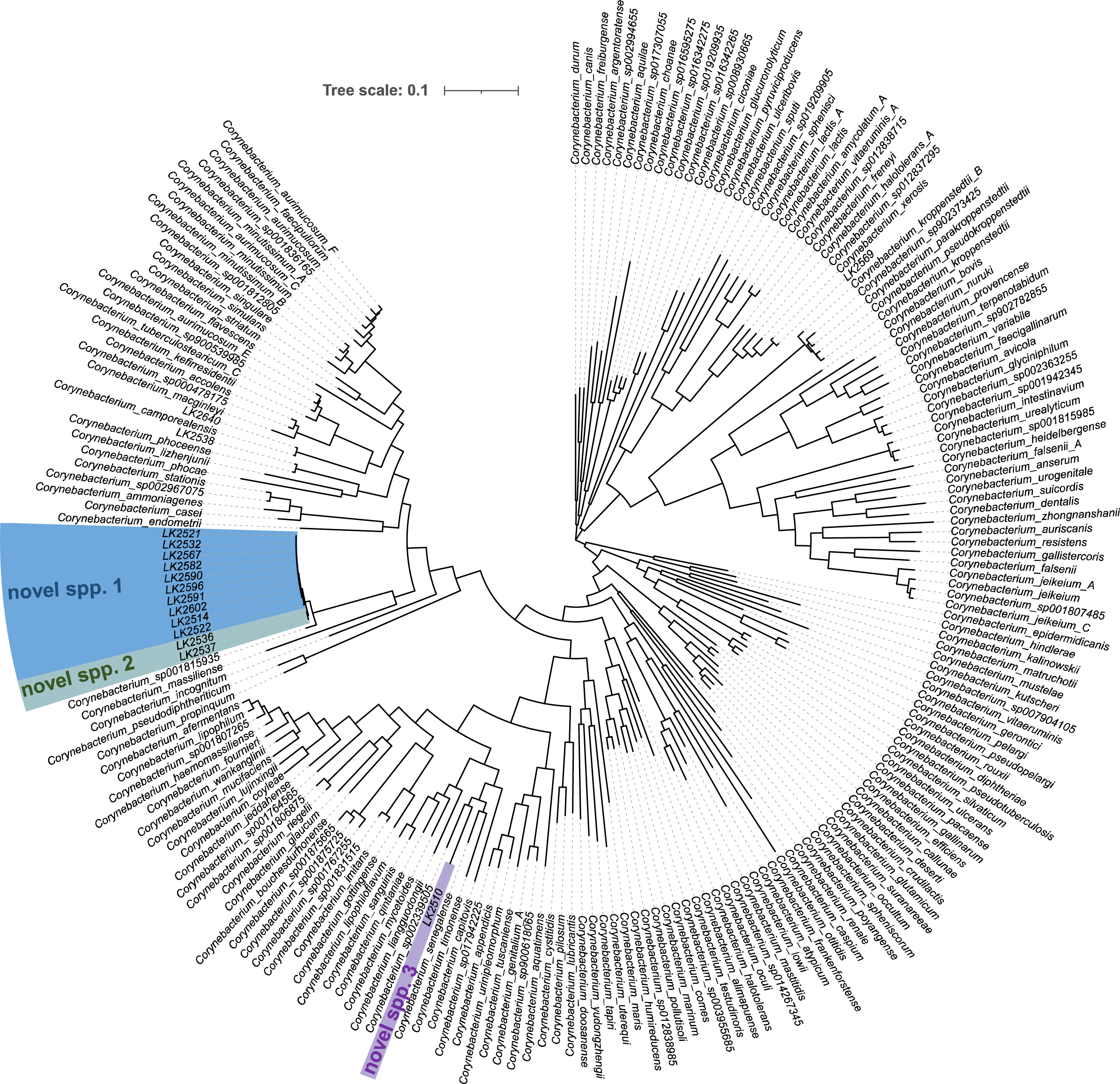
**Three novel species of *Corynebacterium* were isolated from the porcine skin**. Genomic DNA from the isolates were taxonomically assigned using GTDBk and GTDB vR207. The tree was created using iTOL to show the relatedness of known species of *Corynebacterium* against the ones that we isolated from the porcine skin. Three novel species of *Corynebacterium* were identified and are indicated in blue, green, and purple respectively.

**Figure 4.**
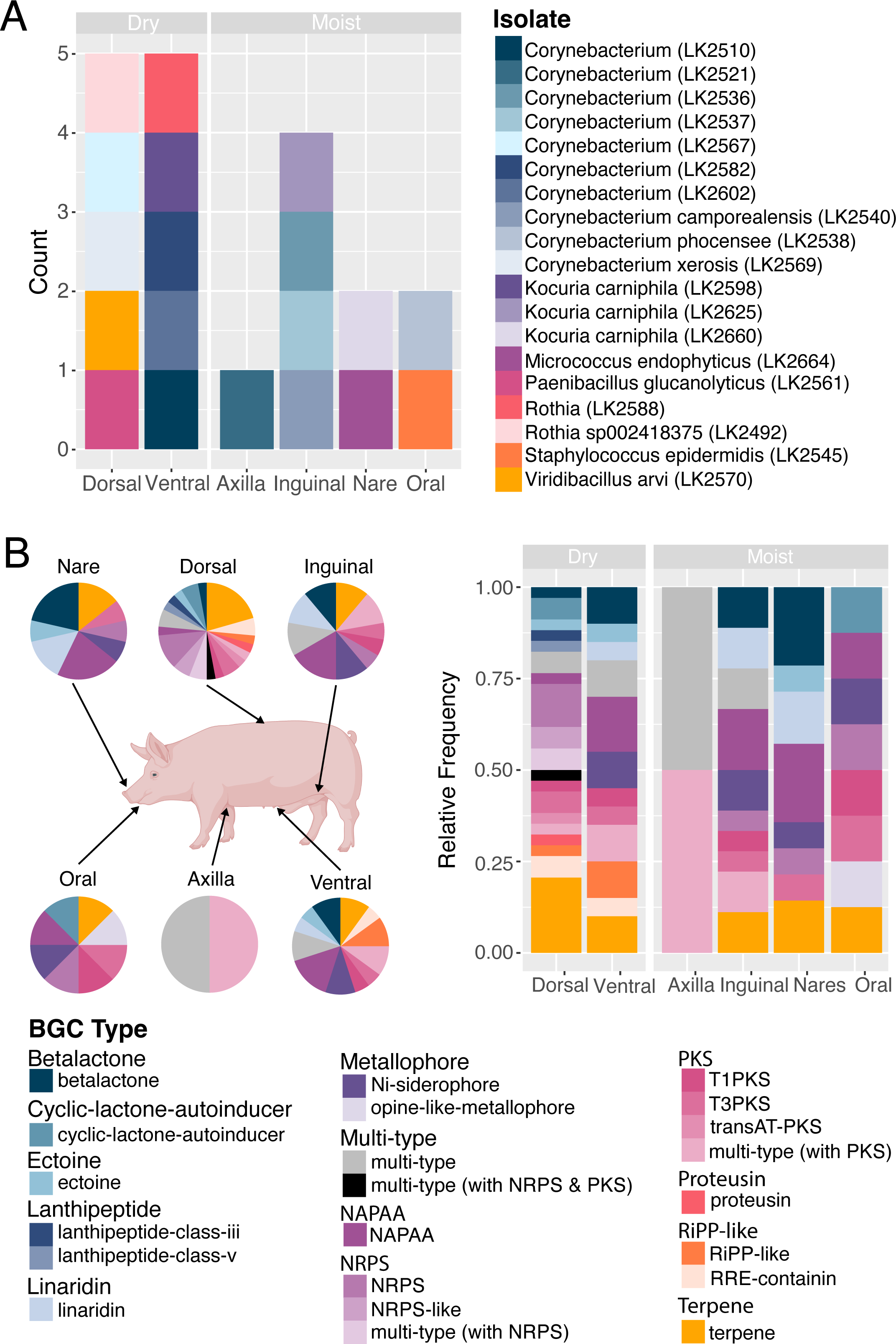
Bacterial isolates from porcine skin encode diverse biosynthetic gene clusters (BGCs). A: Number of isolates from each body site what underwent whole genome sequencing for evaluation of their biosynthetic gene clusters. Body sites grouped by whether they are considered moist or dry sites. Color indicates genera of the isolate. **B:** BGCs within isolate genomes were identified via antiSMASH v7.0. Diagrams illustrate the relative frequencies of types of BGCs found at each respective body site. Details for the BGCs within each individual isolate are displayed in **Supplemental Figure S3 and Supplementary Table S3.**

Next, we examined whether BGCs belonging to the four newly discovered species were novel in comparison to previously cataloged BGCs within BiG-FAM, a database of known BGCs **(Table S3).** The *Paenibacillus glucanolyticus* isolate demonstrated strong antagonism towards fungal pathogens, including *C. albicans* and *A. flavus*, as well as gram-positive bacterial pathogens (**Fig. 2C, Table S2)**. This isolate’s genome featured 15 BGCs, the greatest number of BGCs of the 25 isolates (**Fig. S3, Table S3**). Comparative analysis of BGC predictions with BiG-FAM further revealed that the genome encoded six BGCs with substantial differences from known GCFs. Isolates belonging to two of the novel *Corynebacterium* species, sp. 1 and sp. 2, encoded two BGCs each, both multi-type BGCs. One BGC region was composed of a non-alpha poly-amino acids (NAPAA) and terpene synthesis components while the other BGC region was predicted to encode type 1 polyketide synthase (PKS) and prodigiosin-associated genes. Both BGC types were highly distinct from previously cataloged GCFs. The third novel Corynebacterium species, sp. 3, genome encoded four BGCs. One of these BGCs included a type 1 polyketide synthase, determined to be distinct from known GCFs. The single, novel *Rothia* species also encoded four BGCs within its genome, with only one determined to be distinct from known GCFs. This particular BGC was detected due to the presence of a ribosomally-synthesized and post-translationally modified peptide [RiPP] recognition element. Further investigations into these BGCs that are distinct from known GCFs may uncover novel mechanisms these bacteria utilize to inhibit bacteria and fungi in their surrounding community.

## 4. Discussion

Antimicrobial-resistant infections, particularly those due to antimicrobial-resistant fungi, are on the rise globally (Hendrickson et al., 2019). Human and animal microbiomes are known to provide colonization resistance to potential pathogens, although these mechanisms are largely still under investigation. Characterizing the microbe-microbe interactions that naturally occur within complex microbial communities to maintain community balance is one avenue to identify new antimicrobials. Here, we explored porcine microbiomes for potential antagonistic bacterial-bacterial and bacterial-fungal interactions. Within this work we; i) characterized the bacterial and fungal biomes of porcine skin and feces; ii) screened for bacteria that inhibit the growth of important human pathogens; iii) cultivated and identified four novel species within the *Corynebacterium* and *Rothia* genera with antifungal activity; and iv) identified uncharacterized BGCs that may contribute to these antimicrobial properties. To our knowledge, we are the first to characterize porcine skin fungal communities as well as the first to interrogate the microbial-microbial interactions within these communities. These findings demonstrate how unraveling naturally occurring bacterial-fungal interactions within porcine microbial communities could lead to the discovery of new antimicrobials.

Characterizing fungi from diverse animal-associated environments, including the skin, is still a relatively young pursuit (Tiew et al., 2020). We found that fungal communities were less diverse than their corresponding bacterial communities (**Fig. 1D,H**). The porcine skin mycobiome was dominated by taxa within the *Basidomycota* and *Ascomycota* phyla and in the *Cutaneotrichosporon*, *Apiotrichum* and *Fusarium* genera (**Fig. 1A-B**). Of all the fungal taxa identified, only *Malassezia*, *Aspergillus*, and *Candida* appeared to be to be shared with known human skin communities (Findley et al., 2013; Nguyen and Kalan, 2022). It’s possible that this comparatively low fungal alpha diversity (**Fig. 1D**), greater inter-animal variability (**Fig. 1A-C**), and lower abundance of human associated fungal pathogens (e.g *Candida* species; **Fig. 1B**), are partially due to antifungal molecules produced by the bacteria within the porcine microbiome (**Fig. 2C-D**, **Fig. 4**). Porcine fungal communities are also more similar to those of other mammals, including canines and non-human primates (Fedullo et al., 2013; Meason-Smith et al., 2015). For instance, canine skin communities largely contain *Alternaria*, *Cladosporium*, and *Epicoccum* fungi, all of which were similarly identified in our porcine subjects (**Fig. 1A-B)** (Meason-Smith et al., 2015).

Consistent with previous works characterizing porcine skin microbial communities (Wareham- Mathiassen et al., 2023; Wei et al., 2023), bacterial communities are dominated by taxa within the Bacillota phyla and *Aerococcus* and *Streptococcus* genera (**Fig. 1E-F**). Other shared prominent skin taxa include *Prevotella* and *Lactobacillus* (**Fig. 1E-F)** (Wei et al., 2023). At the genera level, these communities also share key similarities to human skin microbial communities (Byrd et al., 2018; Townsend and Kalan, 2023; Wareham-Mathiassen et al., 2023). As much as 97% of taxa found on pig skin are also found on humans (Wareham-Mathiassen et al., 2023), with both skin communities containing significant proportions of *Corynebacterium*, *Streptococcus*, *Micrococcus*, *Staphylococcus* (**Fig. 1E-F**). However, one notable difference is the absence of *Cutibacterium* (formerly *Propionibacterium*) on porcine skin (**Fig. 1E-F**). The structural and functional similarities between porcine and human skin have long made porcine skin a valuable model for cutaneous and wound healing research (Seaton et al., 2015). Collectively, the microbiome similarities reinforced here and by others (Summerfield et al., 2015; Wareham-Mathiassen et al., 2023), further underscore how porcine skin microbial communities may provide translatable insight into the features that protect humans from pathogens.

*Corynebacterium* species are ubiquitous in the environment (Oliveira et al., 2017) and are frequently found on the skin, nasal, and oral cavity of humans (Esberg et al., 2020; Jensen et al., 2023; Stubbendieck et al., 2019; Townsend and Kalan, 2023) and other mammals (Frischmann et al., 2012; Tang et al., 2020; Wei et al., 2023). Their presence on human and mammalian skin is generally regarded as beneficial and rarely associated with disease (Ahmed et al., 2023; Y. E. Chen et al., 2018; Salamzade et al., 2023b; Swaney et al., 2022; Tang et al., 2020; Wei et al., 2023). Here, we successfully cultured and identified 12 closely related isolates that belong to three novel *Corynebacterium* species (**Fig. 3**). In tandem with previous investigations into the porcine skin microbiome (Wei et al., 2023), isolation of these novel taxa reinforces that *Corynebacterium* are common mammalian skin commensals. Although there are striking similarities between porcine and human skin microbiota at the genus level, these findings also highlight that at the species level porcine and human skin microbial ecosystems are likely distinct (Wei et al., 2023). There is growing interest in understanding the mechanisms *Corynebacterium* utilize to inhibit potential pathogens and modulate local microbial communities. Recent works have reported *C. accolens, C. pseudodiphtheriticum* and other *Corynebacterium* species to inhibit and shift *Staphylococcus aureus,* including methicillin-resistant *S. aureus* (MRSA), into a less virulent state (Hardy et al., 2019; Huang et al., 2022; Menberu et al., 2021; Ramsey et al., 2016; Stubbendieck et al., 2019). In line with these reports several of our *Corynebacterium* isolates, particularly LK2580, LK2599, and LK2632 display strong inhibition of both *S. aureus* or *S. epidermidis* (**Fig. 2, Table S2**). Isolates from *Corynebacterium* novel species 1, LK2521 and LK2567 also displayed moderately strong inhibition of *S. aureus*.

Several *Corynebacterium* isolates also strongly inhibited of multiple fungal pathogens *in vitro* (**Fig 2, Table S2**). This includes, *C. phoceense* (LK2538), *C*. *camporealensis* (LK2640), and several *Corynebacterium* novel species (e.g. LK2521, LK2567, LK2582). Previous work has found *C. amycolatum,* a common human commensal to inhibit the growth and biofilm formation of *Candida* species, including *C. albicans* (Gladysheva et al., 2023)*. Corynebacterium* species from the Gulf of Mannar were also noted to inhibit plant pathogens *Aspergillus niger* and *Alternaria alternata* (Dhinakaran et al., 2012). *Corynebacterium xerosis*, has also been noted to display antagonistic activity against *Candida albicans* as well as *Escherichia coli* (El-Banna, 2006). *C. xerosis* (LK2569) isolated from one of our porcine subjects similarly displayed slight inhibition of *Candida* spp. and strong inhibition of *Cryptococcus neoformans.* Collectively, these antagonistic interactions highlight how *Corynebacterium* may serve as a reservoir of novel antifungals. Select isolates with antimicrobial activity were evaluated via whole genome sequencing (**Fig. 4 and S3**). *Corynebacterium* isolates either contained non-alpha poly-amino acids (NAPAA), Type I PKS, or terpene BGCs. These types of molecules have a range of bioactivities (Esmaeel et al., 2018; Ishaque et al., 2020; Kallscheuer et al., 2019)1/12/2024 6:52:00 AM. These molecules play critical roles in modulating microbial communities through facilitating quorum-sensing and microbe-microbe communications, sequestering nutrients, or directly inhibiting the growth of neighboring bacteria or fungi.

*Rothia* are prevalent members of the upper airway, oral, gut, and skin microbiota in pigs and humans (Akomoneh et al., 2023; Oliveira et al., 2022; Strube et al., 2018; Stubbendieck et al., 2023).We successfully isolated *Rothia* sp002418375 as well as a novel *Rothia* species (**Table S1**). Both isolates displayed strong inhibition of fungal pathogens *in vitro*, with *Rothia* novel species 1 (LK2588) displaying strong inhibition against *Cryptococcus neoformans* and *Candida albicans* and *Rothia* sp002418375 (LK4492) showing significant inhibition of nearly all the fungi within the pathogen panel (**Fig. 2, Table S2**). Multiple sources have reported *Rothia* isolates to encode diverse BGCs (e.g. non-ribosomal peptide synthases [NRPSs], polyketidesynthases [PKSs], and ribosomally-synthesized and post-translationally modified peptides [RiPPs]) ,which can produce antibiotic peptides siderophores and other secondary metabolites capable of modulating the microbiome (Akomoneh et al., 2023; Oliveira et al., 2022). In line with these findings, the porcine skin associated *Rothia* isolates contain multiple BGCs including RIPPs, NAPAA, and terpenes (**Fig. S3**).

We also found *Paenibacillus glucanolyticus* (LK2561) isolated from the dorsal porcine skin to display significant inhibition of several fungal and gram-positive bacterial pathogens (**Fig. 2, Table S2**). This *P. glucanolyticus* encoded 15 distinct BGC’s (**Fig. S3**), consistent with previous reports of other P*aenibacillus* species encoding diverse BGCs (Lebedeva et al., 2021). One notable BGC identified was an 82 kB *trans*-acyltransferase polyketide synthase (*trans*-AT PKS). *Trans*-AT PKSs are involved in the biosynthesis of natural products (Helfrich and Piel, 2016), which can hold antifungal properties (Matilla et al., 2015). Collectively, the BGCs identified within *Paenibicillus, Rothia, Corynebacterium,* highlighted above, as well as those within several other isolated bacteria (**Fig. 4, S3**) underscore the potential breadth of bacterially produced molecules to inhibit the colonization and overgrowth of fungi within these communities (**Fig. 2C- D**).

Limitations of this study include a small number of animals from a single research center, limiting the determination of variability between multiple animals and body site variability. Swine, like all animals, regularly acquire transient microbes from their environment and there is always a possibility that a microbe isolated via culture or detected via DNA sequencing is a transient member of the microbiome. Finally, the bioassays are also intentionally designed to screen for interactions in a pairwise manner under *in vitro* settings. Future works will aim to validate these antagonistic interactions in more complex environments mimicking the skin microbiome diversity and ecosystem.

Fungal infections represent a silent epidemic, often going undiagnosed, leading to poor quality data and estimates of true burden. More, a lack in the supply of effective antifungals is not confined to the healthcare space. Fungal plant pathogens threaten the agriculture industry and entire ecosystems. The discovery of novel species of bacteria producing antifungal molecules in a common and well-studied animal such as the domestic pig is exciting and suggests there is much more biological and chemical diversity left to discover. Given the similarity in skin physiology, architecture, and microbiome of pigs and humans, this work provides the foundation for the discovery of new and safe antimicrobial therapeutics selected for by a mammalian host while addressing fundamental gaps in our understanding of how bacteria and fungi interact.

## Acknowledgements

Special thanks to the University of Wisconsin Biotechnology Center for their assistance with DNA extraction and sequencing.

## CRediT Authorship Contribution Statement

**Karinda F. De La Cruz:** formal analysis, investigation, methodology, visualization, writing – draft, reviewing & editing. **Elizabeth C. Townsend:** formal analysis, investigation, methodology, visualization, writing – draft, reviewing & editing. **JZ Alex Cheong:** formal analysis, investigation, writing – draft, reviewing & editing. **Rauf Salamzade:** investigation, methodology, visualization, writing – draft, reviewing & editing. **Aiping Liu:** investigation, writing – reviewing & editing. **Shelby Sandstrom:** investigation, methodology, visualization, formal analysis, writing – reviewing & editing. **Evelin Davila:** investigation, visualization, writing – draft. **Lynda Huang:** investigation, writing – reviewing & editing. **Kayla H. Xu:** investigation, writing – reviewing & editing. **Sherrie Y. Wu:** investigation, writing – reviewing & editing. **Jennifer J. Meudt:** resources, writing – reviewing & editing. **Dhanansayan Shanmuganayagam:** resources, writing – reviewing & editing. **Angela L.F. Gibson:** conceptualization, investigation, supervision, writing – reviewing & editing. **Lindsay R. Kalan:** conceptualization, methodology, investigation, resources, supervision, writing – draft, reviewing & editing, funding acquisition.

## Supplemental Figure Legends

**Supplemental Figure S1. Bioassay protocol and scoring examples. A:** Visual depiction of the bioassay protocol utilized to evaluate the antimicrobial activity of isolates against a panel of human pathogens. **B**: Example of a well with a score of zero, indicating that the isolate did not inhibit pathogen growth. **C**: Example of a well with a score of one, denoting that the isolate slowed pathogen growth. **D:** Example of a well with a score of two, marking significantly reduced pathogen growth. **E:** Example of a well with an inhibition score of three, indicating complete inhibition of pathogen growth.

**Supplemental Figure S2. Differential abundance of the fungi and bacteria on the porcine skin.** Differential relative abundance of fungal and bacterial taxa between porcine subjects or across skin sites was evaluated with MaAsLin2. For all plots, the false discovery rate (FDR) corrected p-value with the Benjamini-Hochberg correction as well as the coefficient quantifying the difference in the relative abundance between the groups are indicated. **A:** Ascomycota are significantly more abundant in the fecal fungal communities of porcine subject. **B-D:** Differential relative abundance of bacterial taxa at the genera (B-C) and phyla level (D) between the dorsal and ventral skin sites. **E-G:** Differential relative abundance of bacterial taxa within fecal communities at the genera (E-F) and phyla level (G) between each of the animals. **H-I:** Differential relative abundance of bacterial taxa within skin microbial communities at the genera (E-F) and phyla level (G) between each of the animals.

**Supplemental Figure S3. Bacteria isolated from porcine skin encode for diverse biosynthetic gene clusters (BGCs) discovered bacteria isolated found on the porcine skin.** Plot displaying the number of each BGC type within each individual isolate.

## Supplemental Table Legends

**Supplemental Table S1: Taxonomic classifications of bacterial isolates from pig skin identified via whole genome sequencing.**

**Supplemental Table S2. Fractional Inhibition by individual isolates from the porcine skin.** Fractional inhibition was calculated for each isolate to create a normalized score from zero to one for each pathogen group. Zero indicates no inhibition was dedicated within the specified pathogen group. One means complete inhibition against all pathogens in the specified group.

**Supplemental Table S3: Biosynthetic gene clusters (BGCs) within bacteria isolated from porcine skin.** The bacterial isolate, body site the bacteria was isolated from, BGC, product predicted to be produced are indicated.

## References

1. Ahmed, N., Joglekar, P., Deming, C., NISC Comparative Sequencing Program, Lemon, K.P., Kong, H.H., Segre, J.A., Conlan, S., Barnabas, B.B., Black, S., Bouffard, G.G., Brooks, S.Y., Crawford, J., Marfani, H., Dekhtyar, L., Han, J., Ho, S.-L., Legaspi, R., Maduro, Q.L., Masiello, C.A., McDowell, J.C., Montemayor, C., Mullikin, J.C., Park, M., Riebow, N.L., Schandler, K., Schmidt, B., Sison, C., Stantripop, S., Thomas, J.W., Thomas, P.J., Vemulapalli, M., Young, A.C., 2023. Genomic characterization of the *C. tuberculostearicum* species complex, a prominent member of the human skin microbiome. mSystems 8, e00632–23. 10.1128/msystems.00632-23

2. Akomoneh, E.A., Gestels, Z., Abdellati, S., Vereecken, K., Bartholomeeusen, K., Van Den Bossche, D., Kenyon, C., Manoharan-Basil, S.S., 2023. Genome Mining Uncovers NRPS and PKS Clusters in Rothia dentocariosa with Inhibitory Activity against Neisseria Species. Antibiotics 12, 1592. 10.3390/antibiotics12111592

3. Bisanz, J.E., 2018. qiime2R: Importing QIIME2 artifacts and associated data into R sessions. Version 0.99.

4. Blin, K., Shaw, S., Augustijn, H.E., Reitz, Z.L., Biermann, F., Alanjary, M., Fetter, A., Terlouw, B.R., Metcalf, W.W., Helfrich, E.J.N., van Wezel, G.P., Medema, M.H., Weber, T., 2023. antiSMASH 7.0: new and improved predictions for detection, regulation, chemical structures and visualisation. Nucleic Acids Res. 51, W46–W50. 10.1093/nar/gkad344

5. Bolyen, E., Rideout, J.R., Dillon, M.R., Bokulich, N.A., Abnet, C.C., Al-Ghalith, G.A., Alexander, H., Alm, E.J., Arumugam, M., Asnicar, F., Bai, Y., Bisanz, J.E., Bittinger, K., Brejnrod, A., Brislawn, C.J., Brown, C.T., Callahan, B.J., Caraballo-Rodríguez, A.M., Chase, J., Cope, E.K., Da Silva, R., Diener, C., Dorrestein, P.C., Douglas, G.M., Durall, D.M., Duvallet, C., Edwardson, C.F., Ernst, M., Estaki, M., Fouquier, J., Gauglitz, J.M., Gibbons, S.M., Gibson, D.L., Gonzalez, A., Gorlick, K., Guo, J., Hillmann, B., Holmes, S., Holste, H., Huttenhower, C., Huttley, G.A., Janssen, S., Jarmusch, A.K., Jiang, L., Kaehler, B.D., Kang, K.B., Keefe, C.R., Keim, P., Kelley, S.T., Knights, D., Koester, I., Kosciolek, T., Kreps, J., Langille, M.G.I., Lee, J., Ley, R., Liu, Y.- X., Loftfield, E., Lozupone, C., Maher, M., Marotz, C., Martin, B.D., McDonald, D., McIver, L.J., Melnik, A.V., Metcalf, J.L., Morgan, S.C., Morton, J.T., Naimey, A.T., Navas-Molina, J.A., Nothias, L.F., Orchanian, S.B., Pearson, T., Peoples, S.L., Petras, D., Preuss, M.L., Pruesse, E., Rasmussen, L.B., Rivers, A., Robeson, M.S., Rosenthal, P., Segata, N., Shaffer, M., Shiffer, A., Sinha, R., Song, S.J., Spear, J.R., Swafford, A.D., Thompson, L.R., Torres, P.J., Trinh, P., Tripathi, A., Turnbaugh, P.J., Ul-Hasan, S., van der Hooft, J.J.J., Vargas, F., Vázquez-Baeza, Y., Vogtmann, E., von Hippel, M., Walters, W., Wan, Y., Wang, M., Warren, J., Weber, K.C., Williamson, C.H.D., Willis, A.D., Xu, Z.Z., Zaneveld, J.R., Zhang, Y., Zhu, Q., Knight, R., Caporaso, J.G., 2019. Reproducible, interactive, scalable and extensible microbiome data science using QIIME 2. Nat. Biotechnol. 37, 852–857. 10.1038/s41587-019-0209-9

8. Byrd, A.L., Belkaid, Y., Segre, J.A., 2018. The human skin microbiome. Nat. Rev. Microbiol. 16, 143–155. 10.1038/nrmicro.2017.157

9. Callahan, B.J., McMurdie, P.J., Rosen, M.J., Han, A.W., Johnson, A.J.A., Holmes, S.P., 2016. DADA2: High-resolution sample inference from Illumina amplicon data. Nat. Methods 13, 581–583. 10.1038/nmeth.3869

10. Chaumeil, P.-A., Mussig, A.J., Hugenholtz, P., Parks, D.H., 2020. GTDB-Tk: a toolkit to classify genomes with the Genome Taxonomy Database. Bioinformatics 36, 1925–1927. 10.1093/bioinformatics/btz848

11. Chen, S., Zhou, Y., Chen, Y., Gu, J., 2018. fastp: an ultra-fast all-in-one FASTQ preprocessor. Bioinformatics 34, i884–i890. 10.1093/bioinformatics/bty560

12. Chen, Y.E., Fischbach, M.A., Belkaid, Y., 2018. Skin microbiota–host interactions. Nature 553, 427–436. 10.1038/nature25177

13. Chklovski, A., Parks, D.H., Woodcroft, B.J., Tyson, G.W., 2023. CheckM2: a rapid, scalable and accurate tool for assessing microbial genome quality using machine learning. Nat. Methods 20, 1203–1212. 10.1038/s41592-023-01940-w

14. Cruz, N., Abernathy, G.A., Dichosa, A.E.K., Kumar, A., 2022. The Age of Next-Generation Therapeutic-Microbe Discovery: Exploiting Microbe-Microbe and Host-Microbe Interactions for Disease Prevention. Infect. Immun. 90, e00589–21. 10.1128/iai.00589-21

15. Davis, N.M., Proctor, D.M., Holmes, S.P., Relman, D.A., Callahan, B.J., 2018. Simple statistical identification and removal of contaminant sequences in marker-gene and metagenomics data. Microbiome 6, 226. 10.1186/s40168-018-0605-2

16. Dhinakaran, A., Rajasekaran, R., Jayalakshmi, S., 2012. Antiphytopathogenic activity of bacterial protein of a marine Corynebacterium sp. isolated from Mandapam, Gulf of Mannar. J. Biopestic. 5, 17–22.

17. El-Banna, N.M., 2006. Effect of carbon source on the antimicrobial activity of Corynebacterium kutscheri and Corynebacterium xerosis. Afr. J. Biotechnol. 5, 833–835.

18. Esberg, A., Barone, A., Eriksson, L., Lif Holgerson, P., Teneberg, S., Johansson, I., 2020. Corynebacterium matruchotii Demography and Adhesion Determinants in the Oral Cavity of Healthy Individuals. Microorganisms 8, 1780. 10.3390/microorganisms8111780

19. Esmaeel, Q., Pupin, M., Jacques, P., Leclère, V., 20180Nonribosomal peptides and polyketides of Burkholderia: new compounds potentially implicated in biocontrol and pharmaceuticals. Environ. Sci. Pollut. Res. 25, 29794–29807. 10.1007/s11356-017-9166-3

20. Fedullo, J.D.L., Rossi, C.N., Gambale, W., Germano, P.M.L., Larsson, C.E., 2013. Skin mycoflora of *C ebus* primates kept in captivity and semicaptivity. J. Med. Primatol. 42, 293–299. 10.1111/jmp.12056

21. Findley, K., Oh, J., Yang, J., Conlan, S., Deming, C., Meyer, J.A., Schoenfeld, D., Nomicos, E., Park, M., Kong, H.H., Segre, J.A., 2013. Topographic diversity of fungal and bacterial communities in human skin. Nature 498, 367–370. 10.1038/nature12171

22. Frischmann, A., Knoll, A., Hilbert, F., Zasada, A.A., Kämpfer, P., Busse, H.-J., 2012. Corynebacterium epidermidicanis sp. nov., isolated from skin of a dog. Int. J. Syst. Evol. Microbiol. 62, 2194–2200. 10.1099/ijs.0.036061-0

23. Galili, T., Jongblets, A., Pilosov, M., 2020. gplots.

22. Gladysheva, I.V., Chertkov, K.L., Cherkasov, S.V., Khlopko, Y.A., Kataev, V.Y., Valyshev, A.V., 2023. Probiotic Potential, Safety Properties, and Antifungal Activities of Corynebacterium amycolatum ICIS 9 and Corynebacterium amycolatum ICIS 53 Strains. Probiotics Antimicrob. Proteins 15, 588–600. 10.1007/s12602-021-09876-3

24. Hardy, B.L., Dickey, S.W., Plaut, R.D., Riggins, D.P., Stibitz, S., Otto, M., Merrell, D.S., 2019. Corynebacterium pseudodiphtheriticum Exploits Staphylococcus aureus Virulence Components in a Novel Polymicrobial Defense Strategy. mBio 10, e02491–18. 10.1128/mBio.02491-18

25. Helfrich, E.J.N., Piel, J., 2016. Biosynthesis of polyketides by trans-AT polyketide synthases. Nat. Prod. Rep. 33, 231–316. 10.1039/C5NP00125K

26. Hendrickson, J.A., Hu, C., Aitken, S.L., Beyda, N., 2019. Antifungal Resistance: a Concerning Trend for the Present and Future. Curr. Infect. Dis. Rep. 21, 47. 10.1007/s11908-019-0702-9

27. Huang, S., Hon, K., Bennett, C., Hu, H., Menberu, M., Wormald, P.-J., Zhao, Y., Vreugde, S., Liu, S., 2022. Corynebacterium accolens inhibits Staphylococcus aureus induced mucosal barrier disruption. Front. Microbiol. 13, 984741. 10.3389/fmicb.2022.984741

28. Ishaque, N.M., Burgsdorf, I., Limlingan Malit, J.J., Saha, S., Teta, R., Ewe, D., Kannabiran, K., Hrouzek, P., Steindler, L., Costantino, V., Saurav, K., 2020. Isolation, Genomic and Metabolomic Characterization of Streptomyces tendae VITAKN with Quorum Sensing Inhibitory Activity from Southern India. Microorganisms 8, 121. 10.3390/microorganisms8010121

29. Jain, C., Rodriguez-R, L.M., Phillippy, A.M., Konstantinidis, K.T., Aluru, S., 2018. High throughput ANI analysis of 90K prokaryotic genomes reveals clear species boundaries. Nat. Commun. 9, 5114. 10.1038/s41467-018-07641-9

30. Jensen, M.G., Svraka, L., Baez, E., Lund, M., Poehlein, A., Brüggemann, H., 2023. Species- and strain-level diversity of Corynebacteria isolated from human facial skin. BMC Microbiol. 23, 366. 10.1186/s12866-023-03129-9

31. Kallscheuer, N., Kage, H., Milke, L., Nett, M., Marienhagen, J., 2019. Microbial synthesis of the type I polyketide 6-methylsalicylate with Corynebacterium glutamicum. Appl. Microbiol. Biotechnol. 103, 9619–9631. 10.1007/s00253-019-10121-9

32. Kautsar, S.A., Blin, K., Shaw, S., Weber, T., Medema, M.H., 2021. BiG-FAM: the biosynthetic gene cluster families database. Nucleic Acids Res. 49, D490–D497. 10.1093/nar/gkaa812

33. Krueger, F., James, F., Ewels, P., Afyounian, E., Weinstein, M., Schuster-Boeckler, B., Hulselmans, G., Sclamons, 2023. FelixKrueger/TrimGalore: v0.6.10 - add default decompression path. 10.5281/ZENODO.7598955

34. Lebedeva, J., Jukneviciute, G., Čepaitė, R., Vickackaite, V., Pranckutė, R., Kuisiene, N., 2021. Genome Mining and Characterization of Biosynthetic Gene Clusters in Two Cave Strains of Paenibacillus sp. Front. Microbiol. 11, 612483. 10.3389/fmicb.2020.612483

35. Lee, M.D., 2019. GToTree: a user-friendly workflow for phylogenomics. Bioinformatics 35, 4162–4164. 10.1093/bioinformatics/btz188

36. Letunic, I., Bork, P., 2019. Interactive Tree Of Life (iTOL) v4: recent updates and new developments. Nucleic Acids Res. 47, W256–W259. 10.1093/nar/gkz239

37. Mallick, H., Rahnavard, A., McIver, L.J., Ma, S., Zhang, Y., Nguyen, L.H., Tickle, T.L., Weingart, G., Ren, B., Schwager, E.H., Chatterjee, S., Thompson, K.N., Wilkinson, J.E., Subramanian, A., Lu, Y., Waldron, L., Paulson, J.N., Franzosa, E.A., Bravo, H.C., Huttenhower, C., 2021. Multivariable association discovery in population-scale meta-omics studies. PLOS Comput. Biol. 17, e1009442. 10.1371/journal.pcbi.1009442

38. Matilla, M.A., Leeper, F.J., Salmond, G.P.C., 2015. Biosynthesis of the antifungal haterumalide, oocydin A , in *S erratia* , and its regulation by quorum sensing, RPOS and HFQ. Environ. Microbiol. 17, 2993–3008. 10.1111/1462-2920.12839

39. McDermott, A., 2022. Drug-resistant fungi on the rise. Proc. Natl. Acad. Sci. 119, e2217948119. 10.1073/pnas.2217948119

40. McMurdie, P.J., Holmes, S., 2013. phyloseq: An R Package for Reproducible Interactive Analysis and Graphics of Microbiome Census Data. PLoS ONE 8, e61217. 10.1371/journal.pone.0061217

41. Meason-Smith, C., Diesel, A., Patterson, A.P., Older, C.E., Mansell, J.M., Suchodolski, J.S., Rodrigues Hoffmann, A., 2015. What is living on your dog’s skin? Characterization of the canine cutaneous mycobiota and fungal dysbiosis in canine allergic dermatitis. FEMS Microbiol. Ecol. 91, fiv139. 10.1093/femsec/fiv139

42. Medema, M.H., Kottmann, R., Yilmaz, P., Cummings, M., Biggins, J.B., Blin, K., De Bruijn, I., Chooi, Y.H., Claesen, J., Coates, R.C., Cruz-Morales, P., Duddela, S., Düsterhus, S., Edwards, D.J., Fewer, D.P., Garg, N., Geiger, C., Gomez-Escribano, J.P., Greule, A., Hadjithomas, M., Haines, A.S., Helfrich, E.J.N., Hillwig, M.L., Ishida, K., Jones, A.C., Jones, C.S., Jungmann, K., Kegler, C., Kim, H.U., Kötter, P., Krug, D., Masschelein, J., Melnik, A.V., Mantovani, S.M., Monroe, E.A., Moore, M., Moss, N., Nützmann, H.-W., Pan, G., Pati, A., Petras, D., Reen, F.J., Rosconi, F., Rui, Z., Tian, Z., Tobias, N.J., Tsunematsu, Y., Wiemann, P., Wyckoff, E., Yan, X., Yim, G., Yu, F., Xie, Y., Aigle, B., Apel, A.K., Balibar, C.J., Balskus, E.P., Barona-Gómez, F., Bechthold, A., Bode, H.B., Borriss, R., Brady, S.F., Brakhage, A.A., Caffrey, P., Cheng, Y.-Q., Clardy, J., Cox, R.J., De Mot, R., Donadio, S., Donia, M.S., Van Der Donk, W.A., Dorrestein, P.C., Doyle, S., Driessen, A.J.M., Ehling-Schulz, M., Entian, K.-D., Fischbach, M.A., Gerwick, L., Gerwick, W.H., Gross, H., Gust, B., Hertweck, C., Höfte, M., Jensen, S.E., Ju, J., Katz, L., Kaysser, L., Klassen, J.L., Keller, N.P., Kormanec, J., Kuipers, O.P., Kuzuyama, T., Kyrpides, N.C., Kwon, H.-J., Lautru, S., Lavigne, R., Lee, C.Y., Linquan, B., Liu, X., Liu, W., Luzhetskyy, A., Mahmud, T., Mast, Y., Méndez, C., Metsä-Ketelä, M., Micklefield, J., Mitchell, D.A., Moore, B.S., Moreira, L.M., Müller, R., Neilan, B.A., Nett, M., Nielsen, J., O’Gara, F., Oikawa, H., Osbourn, A., Osburne, M.S., Ostash, B., Payne, S.M., Pernodet, J.-L., Petricek, M., Piel, J., Ploux, O., Raaijmakers, J.M., Salas, J.A., Schmitt, E.K., Scott, B., Seipke, R.F., Shen, B., Sherman, D.H., Sivonen, K., Smanski, M.J., Sosio, M., Stegmann, E., Süssmuth, R.D., Tahlan, K., Thomas, C.M., Tang, Y., Truman, A.W., Viaud, M., Walton, J.D., Walsh, C.T., Weber, T., Van Wezel, G.P., Wilkinson, B., Willey, J.M., Wohlleben, W., Wright, G.D., Ziemert, N., Zhang, C., Zotchev, S.B., Breitling, R., Takano, E., Glöckner, F.O., 2015. Minimum Information about a Biosynthetic Gene cluster. Nat. Chem. Biol. 11, 625–631. 10.1038/nchembio.1890

43. Menberu, M.A., Liu, S., Cooksley, C., Hayes, A.J., Psaltis, A.J., Wormald, P.-J., Vreugde, S., 2021. Corynebacterium accolens Has Antimicrobial Activity against Staphylococcus aureus and Methicillin-Resistant S. aureus Pathogens Isolated from the Sinonasal Niche of Chronic Rhinosinusitis Patients. Pathogens 10, 207. 10.3390/pathogens10020207

44. Miethke, M., Pieroni, M., Weber, T., Brönstrup, M., Hammann, P., Halby, L., Arimondo, P.B., Glaser, P., Aigle, B., Bode, H.B., Moreira, R., Li, Y., Luzhetskyy, A., Medema, M.H., Pernodet, J.-L., Stadler, M., Tormo, J.R., Genilloud, O., Truman, A.W., Weissman, K.J., Takano, E., Sabatini, S., Stegmann, E., Brötz-Oesterhelt, H., Wohlleben, W., Seemann, M., Empting, M., Hirsch, A.K.H., Loretz, B., Lehr, C.-M., Titz, A., Herrmann, J., Jaeger, T., Alt, S., Hesterkamp, T., Winterhalter, M., Schiefer, A., Pfarr, K., Hoerauf, A., Graz, H., Graz, M., Lindvall, M., Ramurthy, S., Karlén, A., Van Dongen, M., Petkovic, H., Keller, A., Peyrane, F., Donadio, S., Fraisse, L., Piddock, L.J.V., Gilbert, I.H., Moser, H.E., Müller, R., 2021. Towards the sustainable discovery and development of new antibiotics. Nat. Rev. Chem. 5, 726–749. 10.1038/s41570-021-00313-1

45. Murray, C.J.L., Ikuta, K.S., Sharara, F., Swetschinski, L., Robles Aguilar, G., Gray, A., Han, C., Bisignano, C., Rao, P., Wool, E., Johnson, S.C., Browne, A.J., Chipeta, M.G., Fell, F., Hackett, S., Haines-Woodhouse, G., Kashef Hamadani, B.H., Kumaran, E.A.P., McManigal, B., Achalapong, S., Agarwal, R., Akech, S., Albertson, S., Amuasi, J., Andrews, J., Aravkin, A., Ashley, E., Babin, F.-X., Bailey, F., Baker, S., Basnyat, B., Bekker, A., Bender, R., Berkley, J.A., Bethou, A., Bielicki, J., Boonkasidecha, S., Bukosia, J., Carvalheiro, C., Castañeda-Orjuela, C., Chansamouth, V., Chaurasia, S., Chiurchiù, S., Chowdhury, F., Clotaire Donatien, R., Cook, A.J., Cooper, B., Cressey, T.R., Criollo-Mora, E., Cunningham, M., Darboe, S., Day, N.P.J., De Luca, M., Dokova, K., Dramowski, A., Dunachie, S.J., Duong Bich, T., Eckmanns, T., Eibach, D., Emami, A., Feasey, N., Fisher-Pearson, N., Forrest, K., Garcia, C., Garrett, D., Gastmeier, P., Giref, A.Z., Greer, R.C., Gupta, V., Haller, S., Haselbeck, A., Hay, S.I., Holm, M., Hopkins, S., Hsia, Y., Iregbu, K.C., Jacobs, J., Jarovsky, D., Javanmardi, F., Jenney, A.W.J., Khorana, M., Khusuwan, S., Kissoon, N., Kobeissi, E., Kostyanev, T., Krapp, F., Krumkamp, R., Kumar, A., Kyu, H.H., Lim, C., Lim, K., Limmathurotsakul, D., Loftus, M.J., Lunn, M., Ma, J., Manoharan, A., Marks, F., May, J., Mayxay, M., Mturi, N., Munera-Huertas, T., Musicha, P., Musila, L.A., Mussi-Pinhata, M.M., Naidu, R.N., Nakamura, T., Nanavati, R., Nangia, S., Newton, P., Ngoun, C., Novotney, A., Nwakanma, D., Obiero, C.W., Ochoa, T.J., Olivas-Martinez, A., Olliaro, P., Ooko, E., Ortiz-Brizuela, E., Ounchanum, P., Pak, G.D., Paredes, J.L., Peleg, A.Y., Perrone, C., Phe, T., Phommasone, K., Plakkal, N., Ponce-de-Leon, A., Raad, M., Ramdin, T., Rattanavong, S., Riddell, A., Roberts, T., Robotham, J.V., Roca, A., Rosenthal, V.D., Rudd, K.E., Russell, N., Sader, H.S., Saengchan, W., Schnall, J., Scott, J.A.G., Seekaew, S., Sharland, M., Shivamallappa, M., Sifuentes-Osornio, J., Simpson, A.J., Steenkeste, N., Stewardson, A.J., Stoeva, T., Tasak, N., Thaiprakong, A., Thwaites, G., Tigoi, C., Turner, C., Turner, P., Van Doorn, H.R., Velaphi, S., Vongpradith, A., Vongsouvath, M., Vu, H., Walsh, T., Walson, J.L., Waner, S., Wangrangsimakul, T., Wannapinij, P., Wozniak, T., Young Sharma, T.E.M.W., Yu, K.C., Zheng, P., Sartorius, B., Lopez, A.D., Stergachis, A., Moore, C., Dolecek, C., Naghavi, M., 2022. Global burden of bacterial antimicrobial resistance in 2019: a systematic analysis. The Lancet 399, 629–655. 10.1016/S0140-6736(21)02724-0

46. Navarro-Muñoz, J.C., Selem-Mojica, N., Mullowney, M.W., Kautsar, S.A., Tryon, J.H., Parkinson, E.I., De Los Santos, E.L.C., Yeong, M., Cruz-Morales, P., Abubucker, S., Roeters, A., Lokhorst, W., Fernandez-Guerra, A., Cappelini, L.T.D., Goering, A.W., Thomson, R.J., Metcalf, W.W., Kelleher, N.L., Barona-Gomez, F., Medema, M.H., 2020. A computational framework to explore large-scale biosynthetic diversity. Nat. Chem. Biol. 16, 60–68. 10.1038/s41589-019-0400-9

47. Nguyen, U.T., Kalan, L.R., 2022. Forgotten fungi: the importance of the skin mycobiome. Curr. Opin. Microbiol. 70, 102235. 10.1016/j.mib.2022.102235

48. Oksanen, J., Blanchet, F., Friendly, M., Kindt, R., Legendre, P., McGlinn, D., Minchin, P., O’Hara, R.B., Simpson, G., Stevens, M.H.H., Szoecs, E., Wagner, H., 2020. Package “vegan”: Community Ecology Package.

49. Oliveira, A., Oliveira, L.C., Aburjaile, F., Benevides, L., Tiwari, S., Jamal, S.B., Silva, A., Figueiredo, H.C.P., Ghosh, P., Portela, R.W., De Carvalho Azevedo, V.A., Wattam, A.R., 2017. Insight of Genus Corynebacterium: Ascertaining the Role of Pathogenic and Non-pathogenic Species. Front. Microbiol. 8, 1937. 10.3389/fmicb.2017.01937

50. Oliveira, I.M.F.D., Ng, D.Y.K., Van Baarlen, P., Stegger, M., Andersen, P.S., Wells, J.M., 2022. Comparative genomics of Rothia species reveals diversity in novel biosynthetic gene clusters and ecological adaptation to different eukaryotic hosts and host niches. Microb. Genomics 8. 10.1099/mgen.0.000854

51. Olm, M.R., Brown, C.T., Brooks, B., Firek, B., Baker, R., Burstein, D., Soenjoyo, K., Thomas, B.C., Morowitz, M., Banfield, J.F., 2017. Identical bacterial populations colonize premature infant gut, skin, and oral microbiomes and exhibit different in situ growth rates. Genome Res. 27, 601–612. 10.1101/gr.213256.116

52. Olm, M.R., Crits-Christoph, A., Diamond, S., Lavy, A., Matheus Carnevali, P.B., Banfield, J.F., 2020. Consistent Metagenome-Derived Metrics Verify and Delineate Bacterial Species Boundaries. mSystems 5, e00731–19. 10.1128/mSystems.00731-19

53. Parks, D.H., Chuvochina, M., Rinke, C., Mussig, A.J., Chaumeil, P.-A., Hugenholtz, P., 2022. GTDB: an ongoing census of bacterial and archaeal diversity through a phylogenetically consistent, rank normalized and complete genome-based taxonomy. Nucleic Acids Res. 50, D785–D794. 10.1093/nar/gkab776

54. Ramsey, M.M., Freire, M.O., Gabrilska, R.A., Rumbaugh, K.P., Lemon, K.P., 2016. Staphylococcus aureus Shifts toward Commensalism in Response to Corynebacterium Species. Front. Microbiol. 7. 10.3389/fmicb.2016.01230

55. Saheb Kashaf, S., Proctor, D.M., Deming, C., Saary, P., Hölzer, M., NISC Comparative Sequencing Program, Mullikin, J., Thomas, J., Young, A., Bouffard, G., Barnabas, B., Brooks, S., Han, J., Ho, S., Kim, J., Legaspi, R., Maduro, Q., Marfani, H., Montemayor, C., Riebow, N., Schandler, K., Schmidt, B., Sison, C., Stantripop, M., Black, S., Dekhtyar, M., Masiello, C., McDowell, J., Park, M., Thomas, P., Vemulapalli, M., Taylor, M.E., Kong, H.H., Segre, J.A., Almeida, A., Finn, R.D., 2021. Integrating cultivation and metagenomics for a multi-kingdom view of skin microbiome diversity and functions. Nat. Microbiol. 7, 169–179. 10.1038/s41564-021-01011-w

56. Salamzade, R., Cheong, J.Z.A., Sandstrom, S., Swaney, M.H., Stubbendieck, R.M., Starr, N.L., Currie, C.R., Singh, A.M., Kalan, L.R., 2023a. Evolutionary investigations of the biosynthetic diversity in the skin microbiome using lsaBGC. Microb. Genomics 9. 10.1099/mgen.0.000988

57. Salamzade, R., Manson, A.L., Walker, B.J., Brennan-Krohn, T., Worby, C.J., Ma, P., He, L.L., Shea, T.P., Qu, J., Chapman, S.B., Howe, W., Young, S.K., Wurster, J.I., Delaney, M.L., Kanjilal, S., Onderdonk, A.B., Bittencourt, C.E., Gussin, G.M., Kim, D., Peterson, E.M., Ferraro, M.J., Hooper, D.C., Shenoy, E.S., Cuomo, C.A., Cosimi, L.A., Huang, S.S., Kirby, J.E., Pierce, V.M., Bhattacharyya, R.P., Earl, A.M., 2022. Inter-species geographic signatures for tracing horizontal gene transfer and long-term persistence of carbapenem resistance. Genome Med. 14, 37. 10.1186/s13073-022-01040-y

58. Salamzade, R., Swaney, M.H., Kalan, L.R., 2023b. Comparative Genomic and Metagenomic Investigations of the Corynebacterium tuberculostearicum Species Complex Reveals Potential Mechanisms Underlying Associations To Skin Health and Disease. Microbiol. Spectr. 11, e03578–22. 10.1128/spectrum.03578-22

59. Seaton, M., Hocking, A., Gibran, N.S., 2015. Porcine Models of Cutaneous Wound Healing. ILAR J. 56, 127–138. 10.1093/ilar/ilv016

60. Strube, M.L., Hansen, J.E., Rasmussen, S., Pedersen, K., 2018. A detailed investigation of the porcine skin and nose microbiome using universal and Staphylococcus specific primers. Sci. Rep. 8, 12751. 10.1038/s41598-018-30689-y

61. Stubbendieck, R.M., Dissanayake, E., Burnham, P.M., Zelasko, S.E., Temkin, M.I., Wisdorf, S.S., Vrtis, R.F., Gern, J.E., Currie, C.R., 2023. *Rothia* from the Human Nose Inhibit *Moraxella catarrhalis* Colonization with a Secreted Peptidoglycan Endopeptidase. mBio 14, e00464–23. 10.1128/mbio.00464-23

62. Stubbendieck, R.M., May, D.S., Chevrette, M.G., Temkin, M.I., Wendt-Pienkowski, E., Cagnazzo, J., Carlson, C.M., Gern, J.E., Currie, C.R., 2019. Competition among Nasal Bacteria Suggests a Role for Siderophore-Mediated Interactions in Shaping the Human Nasal Microbiota. Appl. Environ. Microbiol. 85, e02406–18. 10.1128/AEM.02406-18

63. Summerfield, A., Meurens, F., Ricklin, M.E., 2015. The immunology of the porcine skin and its value as a model for human skin. Mol. Immunol. 66, 14–21. 10.1016/j.molimm.2014.10.023

64. Swaney, M.H., Sandstrom, S., Kalan, L.R., 2022. Cobamide Sharing Is Predicted in the Human Skin Microbiome. mSystems 7, e00677–22. 10.1128/msystems.00677-22

65. Tang, S., Prem, A., Tjokrosurjo, J., Sary, M., Van Bel, M.A., Rodrigues-Hoffmann, A., Kavanagh, M., Wu, G., Van Eden, M.E., Krumbeck, J.A., 2020. The canine skin and ear microbiome: A comprehensive survey of pathogens implicated in canine skin and ear infections using a novel next-generation-sequencing-based assay. Vet. Microbiol. 247, 108764. 10.1016/j.vetmic.2020.108764

66. Tiew, P.Y., Mac Aogain, M., Ali, N.A.B.M., Thng, K.X., Goh, K., Lau, K.J.X., Chotirmall, S.H., 2020. The Mycobiome in Health and Disease: Emerging Concepts, Methodologies and Challenges. Mycopathologia. 10.1007/s11046-019-00413-z

67. Townsend, E.C., Kalan, L.R., 2023. The dynamic balance of the skin microbiome across the lifespan. Biochem. Soc. Trans. 51, 71–86. 10.1042/BST20220216

68. Wang, L., Zhang, Y., Xu, J., Wang, C., Yin, L., Tu, Q., Yang, H., Yin, J., 2023. Biosynthetic Gene Clusters from Swine Gut Microbiome. Microorganisms 11, 434. 10.3390/microorganisms11020434

69. Wareham-Mathiassen, S., Pinto Glenting, V., Bay, L., Allesen-Holm, M., Bengtsson, H., Bjarnsholt, T., 2023. Characterization of pig skin microbiome and appraisal as an in vivo subcutaneous injection model. Lab. Anim. 57, 304–318. 10.1177/00236772221136173

70. Wei, M., Flowers, L., Knight, S.A.B., Zheng, Q., Murga-Garrido, S., Uberoi, A., Pan, J.T.-C., Walsh, J., Schroeder, E., Chu, E.W., Campbell, A., Shin, D., Bradley, C.W., Duran-Struuck, R., Grice, E.A., 2023. Harnessing diversity and antagonism within the pig skin microbiota to identify novel mediators of colonization resistance to methicillin-resistant *Staphylococcus aureus*. mSphere 8, e00177–23. 10.1128/msphere.00177-23

71. Wick, R.R., Judd, L.M., Gorrie, C.L., Holt, K.E., 2017. Unicycler: Resolving bacterial genome assemblies from short and long sequencing reads. PLOS Comput. Biol. 13, e1005595. 10.1371/journal.pcbi.1005595

72. Wickham, H., 2016. ggplot2: Elegant Graphics for Data Analysis. Springer-Verlang N. Y.

73. Xia, L., Miao, Y., Cao, A., Liu, Y., Liu, Z., Sun, X., Xue, Y., Xu, Z., Xun, W., Shen, Q., Zhang, N., Zhang, R., 2022. Biosynthetic gene cluster profiling predicts the positive association between antagonism and phylogeny in Bacillus. Nat. Commun. 13, 1023. 10.1038/s41467-022-28668-z

