## Supplementary figures and images for "The Porcine Skin Microbiome Exhibits Broad Fungal Antagonism"

### Sup. Fig. 2

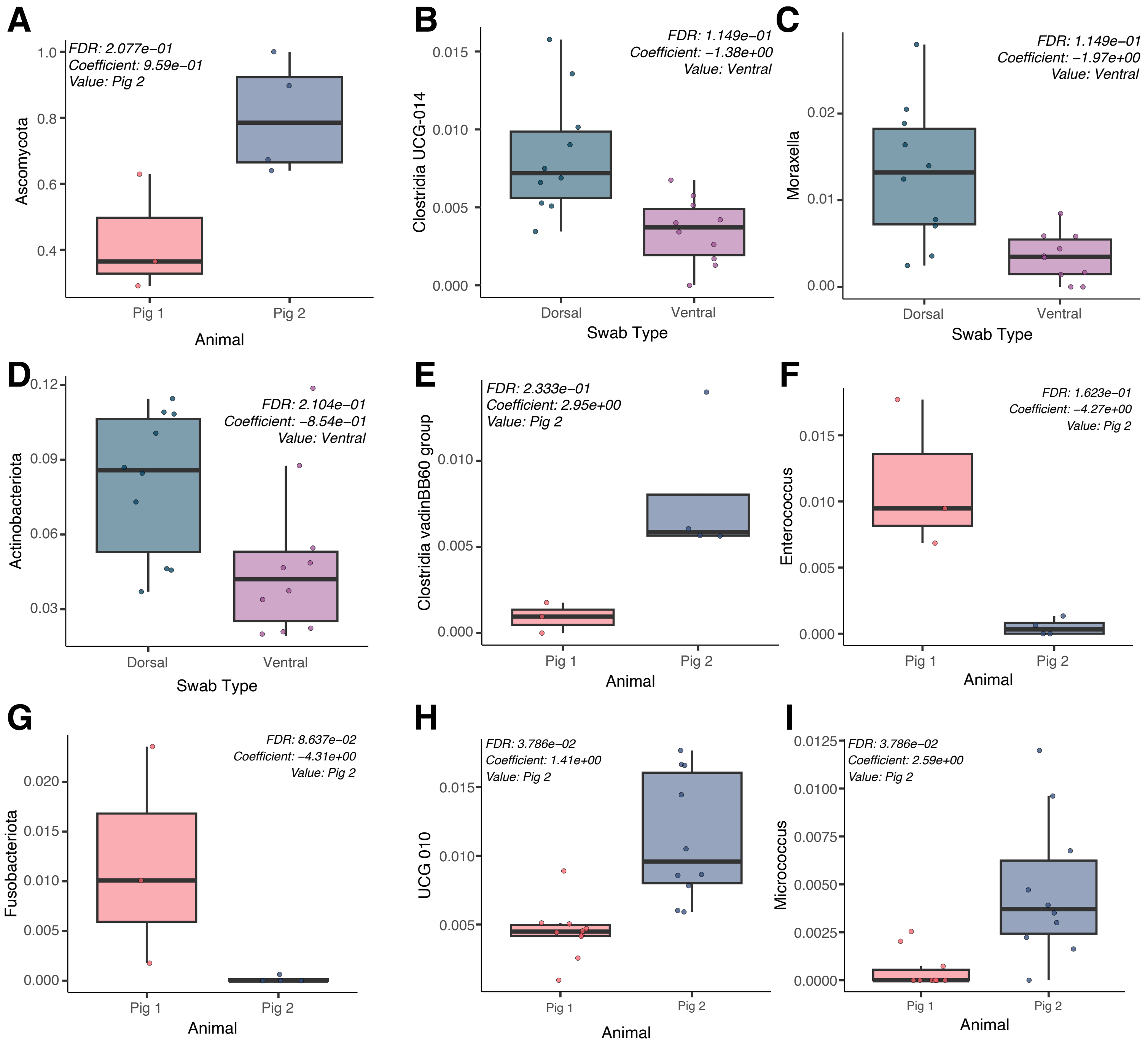

### Sup. Fig. 3

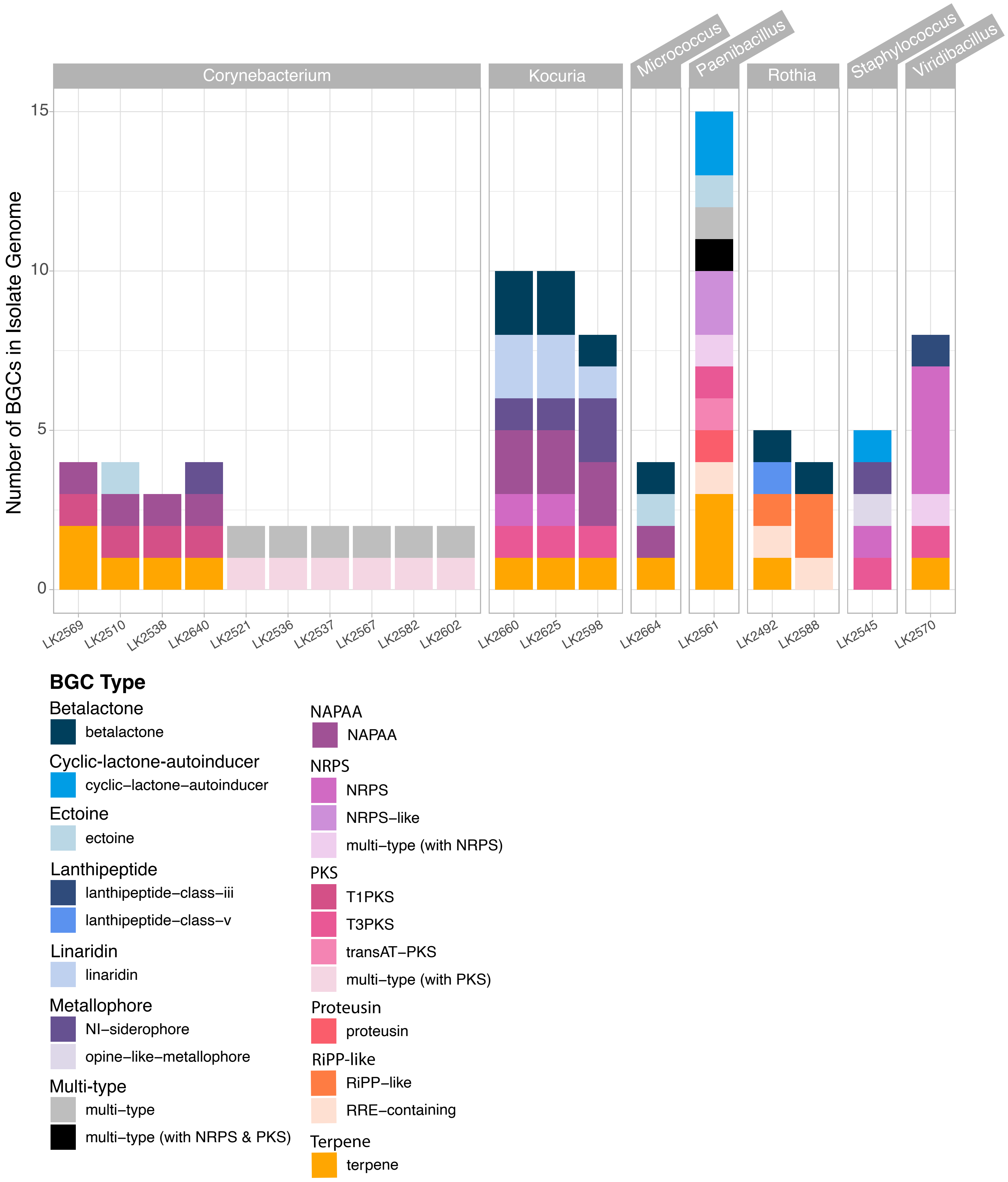

### Sup. Figure 1

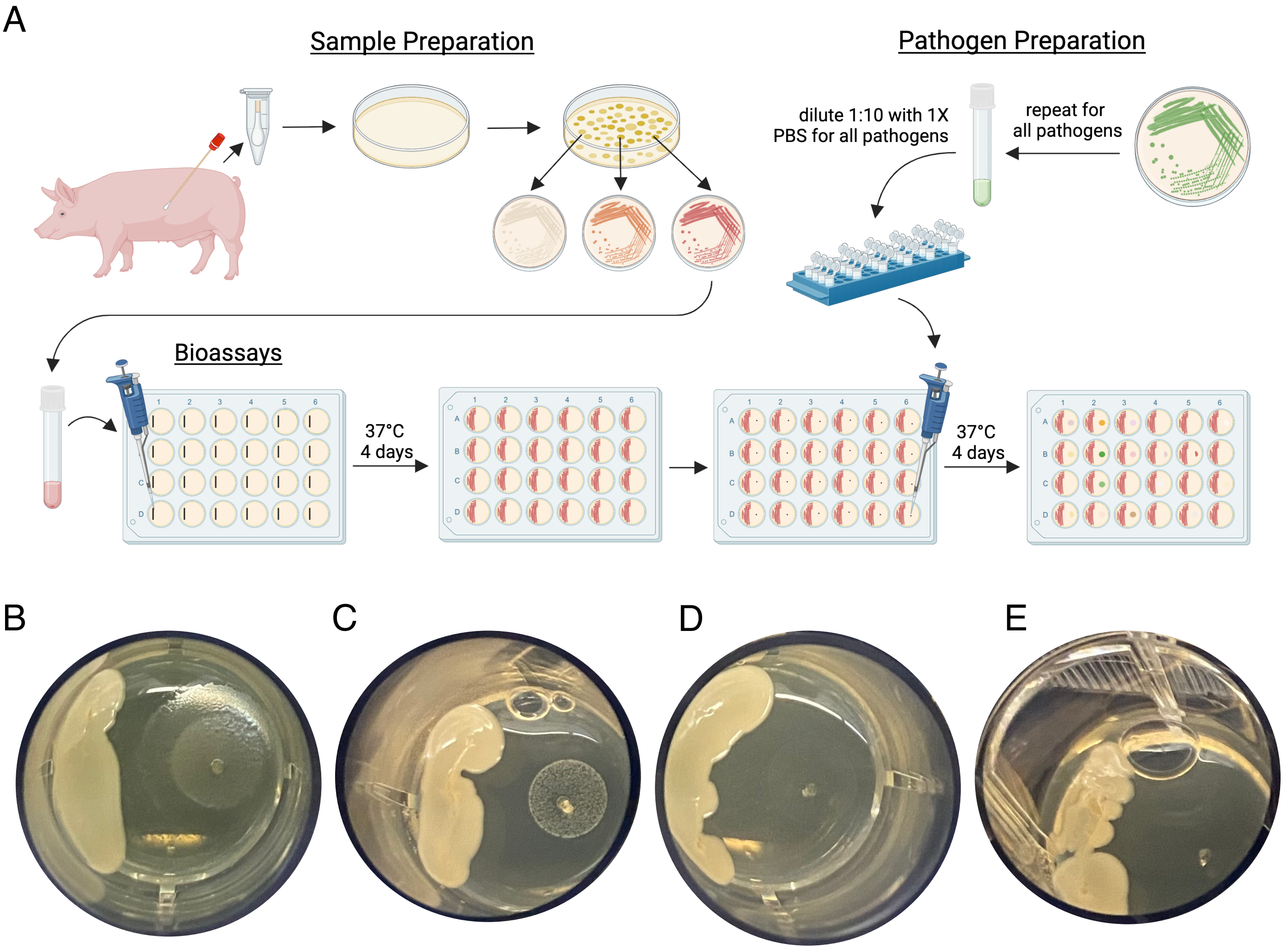
